# Relating visual production and recognition of objects in human visual cortex

**DOI:** 10.1101/724294

**Authors:** Judith E. Fan, Jeffrey D. Wammes, Jordan B. Gunn, Daniel L. K. Yamins, Kenneth A. Norman, Nicholas B. Turk-Browne

## Abstract

Drawing is a powerful tool that can be used to convey rich perceptual information about objects in the world. What are the neural mechanisms that enable us to produce a recognizable drawing of an object, and how does this visual production experience influence how this object is represented in the brain? Here we evaluate the hypothesis that producing and recognizing an object recruit a shared neural representation, such that repeatedly drawing the object can enhance its perceptual discriminability in the brain. We scanned participants using fMRI across three phases of a training study: during training, participants repeatedly drew two objects in an alternating sequence on an MR-compatible tablet; before and after training, they viewed these and two other control objects, allowing us to measure the neural representation of each object in visual cortex. We found that: (1) stimulus-evoked representations of objects in visual cortex are recruited during visually cued production of drawings of these objects, even throughout the period when the object cue is no longer present; (2) the object currently being drawn is prioritized in visual cortex during drawing production, while other repeatedly drawn objects are suppressed; and (3) patterns of connectivity between regions in occipital and parietal cortex supported enhanced decoding of the currently drawn object across the training phase, suggesting a potential substrate for learning how to transform perceptual representations into representational actions. Taken together, our study provides novel insight into the functional relationship between visual production and recognition in the brain.

**Significance Statement:** Humans can produce simple line drawings that capture rich information about their perceptual experiences. However, the mechanisms that support this behavior are not well understood. Here we investigate how regions in visual cortex participate in the recognition of an object and the production of a drawing of it. We find that these regions carry diagnostic information about an object in a similar format both during recognition and production, and that practice drawing an object enhances transmission of information about it to downstream regions. Taken together, our study provides novel insight into the functional relationship between visual production and recognition in the brain.

## Introduction

Although visual cognition is often studied by manipulating externally provided visual information, this ignores our ability to actively control how we engage with our visual environment. For example, people can select which information to encode by shifting their attention (Chun, Golomb, & Turk-Browne, 2011) and can convey which information was encoded by producing a drawing that highlights this information (Bainbridge, Hall, & Baker, 2019; Draschkow, Wolfe, & Vo, 2014). Prior work has provided converging, albeit indirect, evidence that the ability to produce informative visual representations, which we term *visual production*, recruits general-purpose visual processing mechanisms that are also engaged during visual recognition (Fan, Yamins, & Turk-Browne, 2018; James, 2017). The goal of this paper is twofold: first, to more directly characterize the functional role of visual processing mechanisms during visual production; and second, to investigate how repeated visual production influences neural representations that serve perception and action.

With respect to the first goal, our study builds on prior studies that provided evidence for shared computations supporting visual recognition and visual production. For example, recent work has found that activation patterns in human ventral visual stream measured using fMRI (Walther, Chai, Caddigan, Beck, & Li, 2011), as well as activation patterns in higher layers of deep convolutional neural network models of the ventral visual stream (Fan et al., 2018; Yamins et al., 2014), support linear decoding of abstract category information from drawings and color photographs. To what extent are these core visual processing mechanisms also recruited to *produce* a recognizable drawing of those objects? Initial insights bearing on this question have come from human neuroimaging studies investigating the production of handwritten symbols (though not drawings of real-world objects), revealing general engagement of visual regions during both letter production and recognition (Vinci-Booher, Cheng, & James, 2018; James & Gauthier, 2006). However, the format and content of the representations active in these regions during visual production is not yet well understood.

With respect to the second goal, we build on prior work that has investigated the consequences of repeated visual production. In a recent behavioral study, participants who practiced drawing certain objects produced increasingly recognizable drawings and exhibited enhanced perceptual discrimination of morphs of those objects, suggesting that production practice can refine the object representation used for both production and recognition (Fan et al., 2018). These findings resonate with other evidence that visual production can support learning, including maintenance of recently learned information (Peynircioğlu, 1989; Wammes, Meade, & Fernandes, 2016) and enhanced recognition of novel symbols (Longcamp et al., 2008; James & Atwood, 2009; Li & James, 2016). Previous fMRI studies that have investigated the neural mechanisms underlying such learning have found enhanced activation in visual cortex when viewing previously practiced letters (James & Gauthier, 2006; James, 2017), and increased connectivity between visual and parietal regions following handwriting experience (Vinci-Booher, James, & James, 2016). However, these studies have focused on univariate measures of BOLD signal amplitude within regions or when analyzing connectivity, raising the question of whether these changes reflect the recruitment of similar representations across tasks or of co-located but functionally distinct representations for each task.

In the current study, we evaluate the hypothesis that producing and recognizing an object recruit a shared neural representation, such that repeatedly drawing the object can enhance its perceptual discriminability in the brain. Our approach advances prior work that has investigated the neural mechanisms underlying production and recognition in two ways: first, we analyze the *pattern* of activation across voxels to measure the expression and representation of object-specific information; second, we investigate production-related changes to the *organization* of object representations, specifically changes in patterns of voxel-wise connectivity among ventral and dorsal visual regions as a consequence of production practice.

## Materials and Methods

### Participants

Based on initial piloting, we developed a target sample size of 36 participants, across whom all condition and object assignments would be fully counterbalanced. Participants were recruited from the Princeton, NJ community, right-handed, and provided informed consent in accordance with the Princeton IRB. Of the 39 participants who were recruited, 33 participants successfully completed the session. After accounting for technical issues during data acquisition (e.g., excessive head motion), data from 31 participants (11 male, 23.2 years) were retained.

### Stimuli

Four objects from the furniture category were used in this study, based on a prior study (Fan et al., 2018): bed, bench, chair, and table. These objects were represented by 3D mesh models constructed in Autodesk Maya to contain the same number of vertices and the same brown surface texture, and thereby share similar visual properties other than their shape (Fig. 1A). Each of these objects was rendered from a 10°viewing angle (i.e., slightly above) at a fixed distance on a gray background in 40 viewpoints (i.e., each rotated by an additional 9°about the vertical axis).

**Figure 1:**
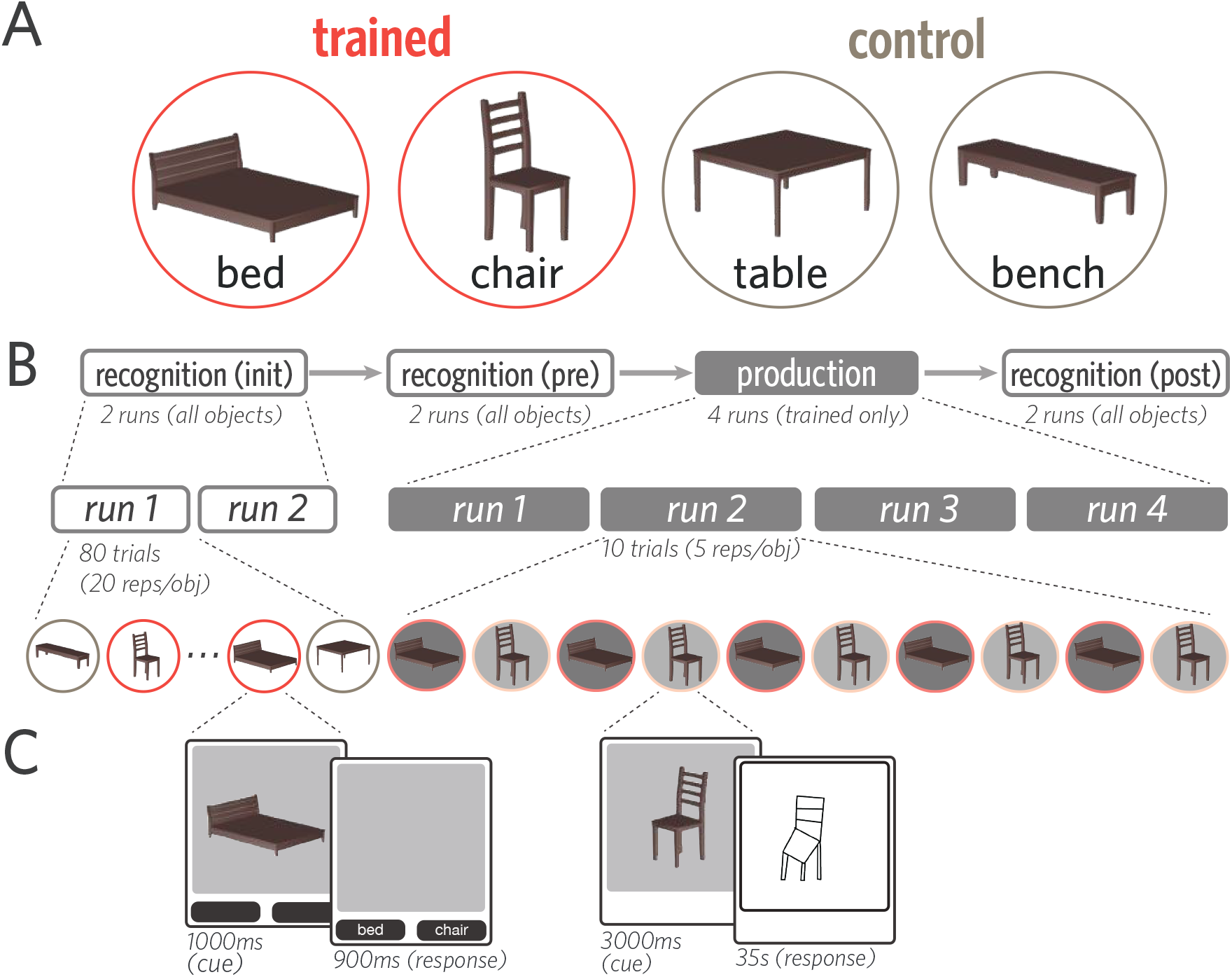
Stimuli, task, and experimental procedure. (A) Four 3D objects were used in this study: bed, bench, chair, and table. Each participant was randomly assigned two of these objects to view and draw repeatedly (trained); the remaining two objects were viewed but never drawn (control). (B) Before and after the production phase, participants viewed all objects while performing a 2AFC recognition task. (C) On each trial of the recognition phase, one of the four objects was briefly presented (1000ms), followed by a 900ms response window. On each trial of the production phase, one trained object was presented (3s), followed by an 35s drawing period (i.e., 23TRs).

### Experimental Design

Each participant was randomly assigned two of the four objects to practice drawing repeatedly (‘trained’ objects). The remaining two objects (‘control’ objects) served as a baseline for changes in neural representations. At the beginning of each session and outside of the scanner, participants were familiarized with each of the four objects while being briefed on the overall experimental procedure. There were four phases in each session (Fig. 1 B&C), all of which were scanned with fMRI: initial recognition (two runs), pre-practice recognition (two runs), production practice (four runs), and a post-practice recognition phase (two runs).

#### Recognition task

Within each of the three recognition phases, participants viewed all four objects in all 40 viewpoints once each and performed an object identification cover task. Repetitions of each object were divided evenly across the two runs of each phase, and in a random order within each run, interleaved with other objects. On each recognition trial, participants were first presented with one of the objects (1000ms). The object then disappeared, and two labels appeared below the image frame, one of which corresponded to the correct object label. Participants then made a speeded forced-choice judgment about which of the two objects they saw by pressing one of two buttons corresponding to each label within a 900ms response window. The assignment of labels to buttons was randomized across trials. Participants did not receive accuracy-related feedback, but received visual feedback if their response was successfully recorded within the response window (selected button highlighted). Inter-stimulus intervals (ISI) were jittered from trial to trial by sampling from the following durations, which appeared in a fixed proportion in each run to ensure equal run lengths: 3000ms ISI (40% trials/run), 4500ms (40%), 6000ms (20%). Each run was 6 minutes in length, and no object appeared in the first or final 12s of each run.

#### Production task

Participants produced drawings on a pressure-sensitive MR-compatible drawing tablet (Hybridmojo) positioned on their lap by using an MR-compatible stylus, which they held like a pencil in the right hand. Before the first drawing run, participants were familiarized with the drawing interface. They practiced producing several closed curves approximately the size of the drawing canvas, to calibrate the extent of drawing movements on the tablet (which they could not directly view) to the appearance of strokes on the canvas. They also practiced drawing two other objects of their choice, providing them with experience drawing more complex shapes using this interface. When participants did not spontaneously generate their own objects to draw, they were prompted to draw a house and a bicycle.

In each of the four runs of the production phase, participants drew both trained objects 5 times each in an alternating order, producing a total of 20 drawings of each object. Each production practice trial had a fixed length of 45s. First, participants were cued with one of the trained objects (3000ms). Following cue offset and a 1000ms delay, a blank drawing canvas of the same dimensions appeared in the same location. Henceforth, we refer to the trained object that is currently being drawn as the *target* object, and to the other trained object not currently being drawn as the *foil* object. Participants then used the subsequent 35s to produce a drawing of the object before the drawing was automatically submitted. Following drawing submission, the canvas was cleared and there was a 6000ms delay until the presentation of the next object cue. Participants were cued with 20 distinct viewpoints of each trained object in a random sequence (18°rotation between neighboring viewpoints), were instructed to to draw each target object in the same orientation as in the image cue, and did not receive performance-related feedback. Each run was 7.7 minutes in length and contained rest periods during the first 12s and final 45s of each run.

### fMRI data acquisition

All fMRI data were collected on a 3T Siemens Skyra scanner with a 64-channel head coil. Functional images were obtained with a multiband echo-planar imaging (EPI) sequence (TR = 1500 ms, TE = 30 ms, flip angle = 70°, acceleration factor = 4, voxel size = 2 mm isotropic), yielding 72 axial slices that provided whole-brain coverage. High resolution T1-weighted anatomical images were acquired with a magnetization-prepared rapid acquisition gradient echo (MPRAGE) sequence (TR = 2530 ms, TE = 3.30 ms, voxel size = 1 mm isotropic, 176 slices, 7°flip angle).

### fMRI data preprocessing

fMRI data were preprocessed with FSL (http://fsl.fmrib.ox.ac.uk). Functional volumes were corrected for slice acquisition time and head motion, high-pass filtered (100s period cutoff), and aligned to the middle volume within each run. For each participant, these individual run-aligned functional volumes were then registered to the anatomical T1 image, using boundary-based registration. All participant-level analyses were performed in participants’ own native anatomical space. For group-level analyses and visualizations, functional volumes were projected into MNI standard space.

### fMRI data analysis

#### Head motion

Given the distal wrist/hand motion required to produce drawings, it was important to measure and verify that there was not extreme head motion during drawing production relative to rest periods (i.e. cue presentation, and delay). For each production run, the time courses for estimated rotations, translations, and absolute and relative displacements, were extracted from the output of MCFLIRT. Functional data were partitioned into production (i.e. the 23 TRs spent drawing in each TR) and rest (i.e., during cue presentation or delay between trials) volumes. We found that there was no difference in rotational movement between production and rest periods (mean = −0.0001; 95% CI = [-0.0003 0.0001]). In fact, there was reliably *less* head movement during production relative to rest, as measured by translation (mean = −0.006; 95% CI = [-0.011 −0.002]), absolute (mean = −0.027; 95% CI = [-0.054 −0.004]) and relative displacement (mean = −0.016; 95% CI = [-0.024 −0.008]).

#### Defining regions of interest in occipitotemporal cortex

We focused our analyses on nine regions of interest (ROIs) in occipitotemporal cortex: V1, V2, lateral occipital cortex (LOC), fusiform (FUS), inferior temporal lobe (IT), parahippocampal cortex (PHC), perirhinal cortex (PRC), entorhinal cortex (EC), and hippocampus (HC). These regions were selected based on prior evidence for their functional involvement in processing. For instance, neurons in V1 and V2 are tuned to the orientation of perceived contours, which constitute simple line drawings and also often define the edges of an object (Hubel & Wiesel, 1968; Gegenfurtner, Kiper, & Fenstemaker, 1996; Kamitani & Tong, 2005; Sayim & Cavanagh, 2011). Likewise, neural populations in higher-level ventral regions, including LOC, FUS, and IT, have been shown to play an important role in representing more abstract invariant properties of objects (Grill-Spector, Kourtzi, & Kanwisher, 2001; Kourtzi & Kanwisher, 2001; Hung, Kreiman, Poggio, & DiCarlo, 2005; Rust & DiCarlo, 2010; Gross, 1992); with medial temporal regions including PHC, PRC, EC, and HC participating in both online visual processing, as well as the formation of visual memories (Murray & Bussey, 1999; Epstein, Graham, & Downing, 2003; Davachi, 2006; Schapiro, Kustner, & Turk-Browne, 2012; Garvert, Dolan, & Behrens, 2017). Masks for each ROI were defined in each participants’ T1 anatomical scan, using FreeSurfer segmentations (http://surfer.nmr.mgh.harvard.edu/).

#### Defining production-related regions in parietal cortex

Motivated by prior work investigating visually-guided action (Vinci-Booher et al., 2018; Goodale & Milner, 1992), we also sought to analyze how sensory information represented in occipital cortex is related to downstream parietal cortex, which is associated with action planning and execution. Accordingly, a parietal lobe ROI mask was also generated for each participant based on their Freesurfer segmentation. To determine which voxels across the whole brain were specifically engaged during production, a group-level univariate activation map was estimated contrasting production vs. rest. To derive these production task-related activation maps, we analyzed each production run with a general linear model (GLM). Regressors were specified for each trained object by convolving a boxcar function, reflecting the total amount of time spent drawing (i.e., 23 TRs, or 34.5 s), with a double-gamma hemodynamic response function (HRF). A univariate contrast was then applied, with equal weighting on the regressors for each trained object, to determine the clusters of voxels that were preferentially active during drawing production, relative to rest. Voxels that exceeded a strict threshold (Z = 3.1) and also lay within the anatomically defined ROI boundaries (in either visual cortex or parietal cortex) were included.

To avoid statistical dependence between this procedure used for voxel selection and for subsequent classifier-based analyses, we defined participant-specific activation maps in a leave-one-participant-out fashion. That is, a held out participant’s production mask was constructed based solely on the basis of task-related activations from all *remaining* participants. Once each participant’s mask was defined, we took the intersection between this map and the participant’s own anatomically defined cortical segmentation to construct the production-related ROIs in V1, V2, LOC and parietal cortex. We had no *a priori* predictions about hemispheric differences, so ROI masks were collapsed over the left and right hemispheres.

#### Measuring object evidence during recognition and production phases

In order to quantify the expression of object-specific information throughout recognition and production, we analyzed the neural activation patterns across voxels associated with each object (Haxby et al., 2001; Kamitani & Tong, 2005; Norman, Polyn, Detre, & Haxby, 2006; Cohen et al., 2017). Specifically, we extracted neural activation patterns evoked by each object cue during recognition, measured 3 TRs following each stimulus offset to account for hemodynamic lag. We used these patterns to train a 4-way logistic regression classifier with L2 regularization to predict the identity of the current object in either held-out recognition data or production data. This procedure was performed separately in each ROI in each participant, and all raw neural activation patterns were z-scored within voxel and within run prior to be used for either classifier training or evaluation.

To measure object evidence during recognition, we applied the classifier in a 2-fold crossvalidated fashion within each of the pre-production and post-production phases, such that for each fold, the data from one run were used as training, while the data from the other run were used for evaluation. Aggregating predictions across folds, we computed the proportion of recognition trials on which the classifier correctly identified the currently viewed object, providing a benchmark estimate of how much object-specific information was available from neural activation patterns during recognition. We constructed 95% confidence intervals (CIs) for estimates of decoding accuracy for each ROI by bootstrap resampling participants 10,000 times.

To measure object evidence during production, we trained the same type of classifier exclusively on data from the initial recognition phase, which minimized statistical dependence on the classifier based on pre- and post-production phases. We then evaluated this classifier on every timepoint while participants produced their drawings, which consisted of the 23 TRs following the offset of the image cue, shifted forward 3 TRs to account for hemodynamic lag (Fig. 2).

**Figure 2:**
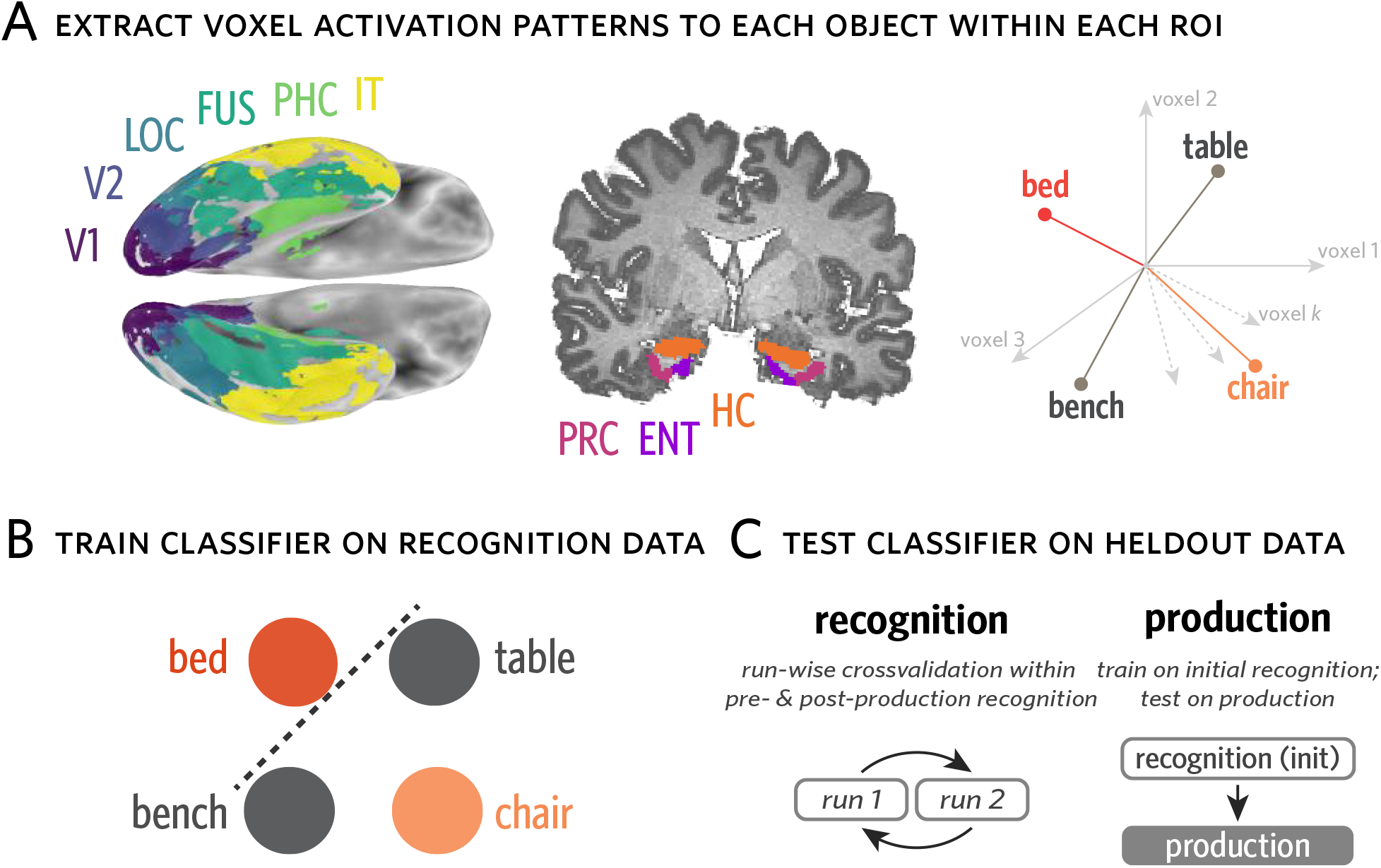
Measuring object evidence in activation patterns during recognition and production. (A) For each participant, anatomical ROIs were defined using FreeSurfer. Activation patterns across voxels in each ROI were extracted for each recognition trial and for all timepoints of each production trial. These activation patterns can be expressed as vectors in a *k*-dimensional vector space, where *k* reflects the number of voxels in a given ROI. (B) Evidence for each object was measured using a 4-way logistic regression classifier trained on activation patterns from recognition runs to predict the current object being viewed or drawn (e.g., bed), and discriminate it from the other three objects (i.e., bench, chair, table). This classifier can be used to measure both the general expression of object-specific information, measured by classification accuracy, as well as the degree of evidence for particular objects, measured by the probabilities it assigns to each. (C) To measure object evidence during recognition, this classifier was trained in a run-wise crossvalidated manner within each of the pre-production and post-production phases. To measure object evidence during production, the same type of classifier was trained on data from the initial recognition phase.

Because this type of classifier assigns a probability value to each object, it can be used to evaluate the strength of evidence for each object at each timepoint. To evaluate the degree to which the currently drawn object (target) was prioritized, we extracted the classifier probabilities assigned to the target, foil, and two control objects on each TR during drawing production. We then used these probabilities to derive metrics that quantify the relative evidence for one object compared to the others. Specifically, we define ‘target selection’ as the log odds ratio between the target and foil objects (ln[*p*(*target*)*/p*(*foil*)]), which captures the degree to which the voxel pattern is more diagnostic of the target than the foil. We define ‘target evidence’ as the log odds ratio between the target and the mean natural-log probabilities assigned to the two control objects for each time point, which captures the degree to which the voxel pattern is more diagnostic of the target than the baseline control objects. We likewise define ‘foil evidence’ as the log odds ratio between the *foil* object and the mean natural log probabilities for the two control objects, which captures the degree to which the voxel pattern is more diagnostic of the *foil* than the baseline control objects. For each ROI within a participant, we compute the average target selection, target evidence, and foil evidence across time points in all four production runs, then aggregate these estimates across participants to compute a group-level estimate for each metric and CI derived via bootstrap resampling of participants 1000 times.

#### Connectivity pattern similarity analysis

The foregoing approach to analyzing multivariate neural representations focused exclusively on spatial activation patterns within anatomically defined regions. However, given that visual production inherently entails the coordination between posterior perceptual and downstream action-oriented systems, we developed an approach to explore how sensory information is transmitted between regions. Specifically, because prior work has indicated that parietal cortex is also engaged during visual production (Vinci-Booher et al., 2018), we measured how activation patterns in visual cortex are related to activation patterns in parietal cortex during drawing production.

For each pair of ROIs (e.g., V1 and Parietal), we extracted the connectivity pattern from every production trial (Fig. 3). Each connectivity pattern consists of the *m* × *n* pairwise temporal correlations between every voxel in one ROI (containing *m* voxels) with every voxel in the second ROI (containing *n* voxels). The temporal correlation between each pair of voxels reflects the correlation between the activation timeseries for the first voxel and the activation timeseries for the second voxel, over all 23 TRs in each production trial.

**Figure 3:**
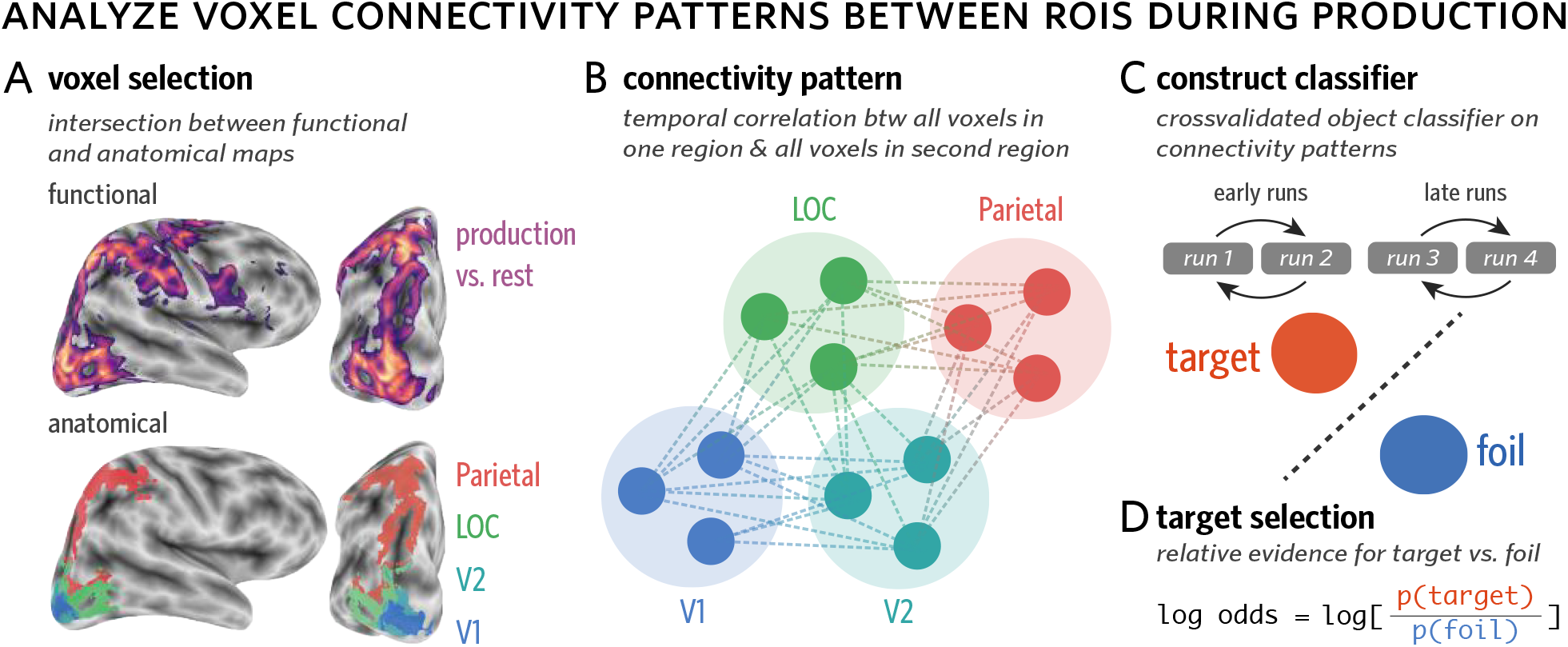
Measuring object evidence in connectivity patterns between regions during production. (A) Voxels in each of several anatomical ROIs (i.e., V1, V2, LOC, Parietal) that were also consistently engaged during the production task were included in this analysis. To determine which voxels were consistently engaged during production, while minimizing statistical dependence between voxel selection and multivoxel pattern analysis, a production task-related activation map was generated in a leave-one-participant-out manner. (B) Connectivity patterns were computed for each trial, for each pair of ROIs. Each connectivity pattern consists of the set of *m* × *n* pairwise temporal correlations between every voxel in one ROI (containing *m* voxels) with every voxel in the second ROI (containing *n* voxels). The temporal correlation between each pair of voxels reflects the correlation between the activation timeseries for the first voxel and the activation timeseries for the second voxel, over all 23 TRs in each production trial. (C) Connectivity patterns were used to construct a 2-way logistic regression classifier to discriminate the currently drawn object (target) from the other trained object (foil). This classifier was trained in a run-wise crossvalidated manner within the first two runs (early) and the final two runs (late) of the production phase. (D) Target selection, the degree to which the target was prioritized over the foil, was defined as the log odds ratio between the target and foil objects.

For each pair of ROIs, we then trained a 2-way logistic regression classifier to discriminate the target vs. foil objects based on these connectivity patterns. The classifier was trained in a run-wise crossvalidated manner within the first two runs (early) and the final two runs (late) of the production phase. To capture the degree to which the connectivity pattern was more diagnostic of the target than the foil, we computed target selection, which was averaged over all trials within a phase (early or late).

Data were fit with a linear mixed-effects regression model (Bates et al., 2015) that included time (early vs. late) as a predictor and random intercepts for different participants. We compared this model to a baseline model that did not include time as a predictor. The reliability of the increase in target selection across time was measured in two ways: first, the model was contrasted with the baseline model, to evaluate the extent to which including time as a predictor improved model fit; second, bootstrapped 95% CIs were computed for each estimate, to evaluate whether they spanned 0 (or chance) and thus determine statistical reliability.

To further evaluate whether the connectivity pattern carried task-related information that was not redundant with the activation patterns within regions, we conducted a control analysis which involved constructing the same type of classifier on the concatenated voxel activation patterns extracted from each ROI, rather than their connectivity pattern.

## Results

### Discriminable object representations in visual cortex during recognition

Following prior work (Haxby et al., 2001; Norman et al., 2006; Cichy, Chen, & Haynes, 2011; Cohen et al., 2017), we hypothesized that there would be consistent information about the identity of each object in visual cortex across repeated presentations during the recognition phase. Specifically, we predicted that the stimulus-evoked pattern of neural activity across voxels in visual cortex upon viewing an object could be used to reliably decode its identity. To test this prediction, we first extracted neural activation patterns evoked by each object during recognition separately for each participant, in each occipitotemporal ROI. We used neural activation patterns extracted from a subset of recognition-phase data to train a 4-way logistic regression classifier that could be used to evaluate decoding accuracy on held-out recognition data in the same regions (Fig. 2). We computed a 2-fold crossvalidated measure of object decoding accuracy (Fig. 4), wherein for each of the pre-production and post-production phases, the 40 repetitions from one of the two runs were used for training the classifier, while the 40 repetitions from the other run were used for evaluation.

**Figure 4:**
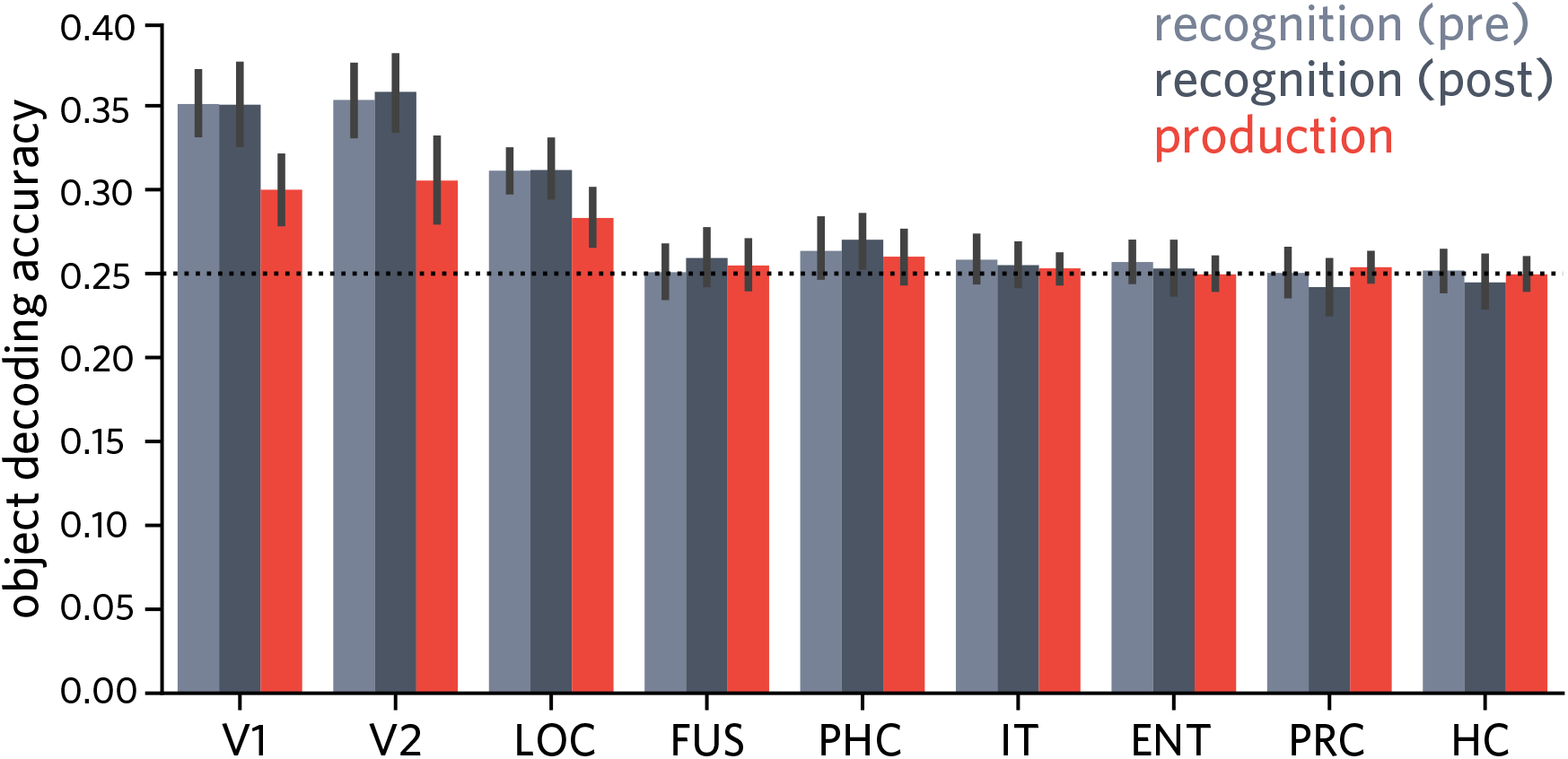
Accuracy of object classifier during pre/post recognition phase and drawing production phase, for each ventral visual region of interest. Error bars reflect 95% CIs.

We found that the identity of the currently viewed object could be reliably decoded in V1, V2, and LOC in the pre recognition phase (95% CIs: V1 = [.332. 370], V2 = [.332. 374], LOC = [.299. 324]; chance=.25; Fig. 4), but not in the more anterior ROIs (95% CIs: FUS = [.236. 266], PHC = [.248. 280], IT = [.245. 272], ENT = [.246. 268], PRC = [.237. 264], HC = [.241. 263]). Likewise, we found that the same early visual regions, as well as PHC, supported above-chance decoding during the post phase (95% CIs: V1 = [.327. 374], V2 = [.337. 379], LOC = [.296. 329], PHC = [.255. 286]), but not the other regions (95% CIs: FUS = [.244. 275], IT = [.242. 268], ENT = [.238. 268], PRC = [.227. 258], HC = [.232. 259]). These results suggest that information about object identity was not uniformly accessible from all regions along the ventral stream, but primarily in occipital cortex, consistent with previous work (Grill-Spector et al., 2001; Güçlü & van Gerven, 2015).

### Similar object representations in visual cortex during recognition and production

The results so far show that there is robust object-specific information evoked by visual recognition of each object in the patterns of neural activity in V1, V2, and LOC. Based on prior work (Fan et al., 2018), we further hypothesized that the neural object representation evoked during recognition would be functionally similar to that recruited during drawing production. Specifically, we predicted that consistency in the patterns of neural activity evoked in visual cortex upon viewing an object could be leveraged to decode the identity of that object during drawing production, even during the period when the object cue was no longer visible. To test this prediction, we evaluated how well a linear classifier trained exclusively on recognition data to decode object identity could generalize to production data in the same regions.

For each ROI in each participant, we used activation patterns evoked by each object across 40 repetitions in two initial recognition runs to train a 4-way logistic regression classifier, which we then applied to each timepoint across the four production practice runs. Critically, we restricted our classifier-based evaluation of production data to the 23 TRs following the offset of the object cue in each trial, providing a measure of the degree to which object-specific information was available in each ROI during production throughout the period when the object was no longer visible. Moreover, we ensured that the data used to train this classifier came from different runs than those used to measure the expression of object-specific information in these regions during the pre- and post-production recognition phases. Averaging over all TRs during production, we found reliable decoding of object identity in V1 (mean = 0.3; 95% CI = [0.280 0.320], chance = .25, Fig. 4), V2 (mean = 0.305; 95% CI = [0.281 0.331]), and LOC (mean = 0.283; 95% CI = [0.267 0.299]), though not in the more anterior ROIs (95% CIs: FUS = [0.241 0.268], PHC = [0.244 0.275], IT = [0.245 0.261], EC = [0.241 0.259], PRC = [0.246 0.262], HC = [0.241 0.258]; Fig. 4).

These results suggest that despite large differences between the two tasks — that is, visual discrimination of a realistic rendering vs. production of a simple sketch based on object information in working memory — there are functional similarities between the visually-evoked representation of objects in occipital cortex (i.e., V1, V2, LOC) and the representation that is recruited during the production of drawings of these objects.

### Sustained selection of target object during production in visual cortex

The findings so far show that the identity of the currently drawn object can be linearly decoded from voxel activation patterns in occipital cortex during drawing production. While this speaks to the overall prioritization of the currently drawn target object in visual cortex, it is unclear whether this prioritization is specific to the target. It may be that both trained objects were activated to a similar and heightened degree during the production phase relative to the control objects, because participants alternated between these objects. On the other hand, this alternation may have led participants to *selectively* prioritize the target object, resulting in the foil object not only being less activated than the target, but also suppressed relative to the control objects. Another question raised by these findings concerned the degree to which object decodability during drawing production was driven by visual recognition of the finished drawing itself, which shared many of the same local visual properties that the object renderings had (e.g., oriented edges), rather than early recruitment of an internal representation of the object that supported drawing production. To tease these possibilities apart, we quantified the relative evidence for each object on every time point during drawing production, in each ventral stream ROI.

We found sustained target evidence (target *>* control) across the production phase in V1 (mean = 0.228; 95% CI = [0.102 0.361]), V2 (mean = 0.227; 95% CI = [0.094 0.360]), and LOC (mean = 0.128; 95% CI = [0.035 0.231]), consistent with the classifier accuracy results reported above. We did not find reliable evidence for sustained target evidence in the other ROIs (95% CIs: FUS = [-0.025 0.222], PHC = [-0.067 0.056], IT = [-0.163 0.026], EC = [-0.113 0.020], PRC = [-0.103 0.018], HC = [-0.047 0.062]).

We also found reliable *negative* foil evidence (foil *<* control) across the production phase again in V1 (mean = −0.449; 95% CI = [-0.601 −0.295]), V2 (mean=−0.481; 95% CI = [-0.701 −0.261]), and LOC (mean = −0.188; 95% CI = [-0.277 −0.095]), suggesting that not only is the task-relevant target object prioritized in these regions, but that the presently task-irrelevant foil object is suppressed. Again, we did not find reliable evidence for sustained foil evidence (in either direction) in the other ROIs (95% CIs: FUS = [-0.170 0.067], PHC = [-0.030 0.072], IT = [-0.154 0.06], EC = [-0.119 0.013], PRC = [-0.05 0.056], HC = [-0.064 0.051]).

Finally, we found sustained target selection (target *>* foil) across the production phase again in V1 (mean = 0.676; 95% CI = [0.449 0.906]), V2 (mean = 0.708; 95% CI = [0.484 0.955]), LOC (mean = 0.316; 95% CI = [0.216 0.423]), and additionally in FUS (mean = 0.151; 95% CI = [0.074 0.229]). Again, we did not find reliable evidence for sustained target selection in the other ventral stream ROIs (95% CIs: PHC = [-0.081 0.0262], IT = [-0.098 0.056], EC = [-0.053 0.063], PRC = [-0.112 0.022], HC = [-0.041 0.068]).

Overall, these results show that the currently drawn object was *selectively* prioritized in occipital cortex, relative to both never-drawn control objects and the other trained object, which was reliably suppressed below the control-object baseline throughout the production phase. Moreover, they reveal reliable target evidence and negative foil evidence at early timepoints in each production trial, prior to drawing completion (Fig. 5B), both of which argue against the account on which accurate object decoding was purely driven by sensory processing of the finished drawing. Taken together, these findings instead suggest the operation of a selection mechanism during drawing production that simultaneously enhances the currently relevant target object representation and suppresses the currently irrelevant foil object representation in early visual cortex, similar to what may be deployed to support selective attention and working memory (Tipper, Weaver, & Houghton, 1994; Serences, Ester, Vogel, & Awh, 2009; Gazzaley & Nobre, 2012; Lewis-Peacock & Postle, 2012; Fan & Turk-Browne, 2013).

**Figure 5:**
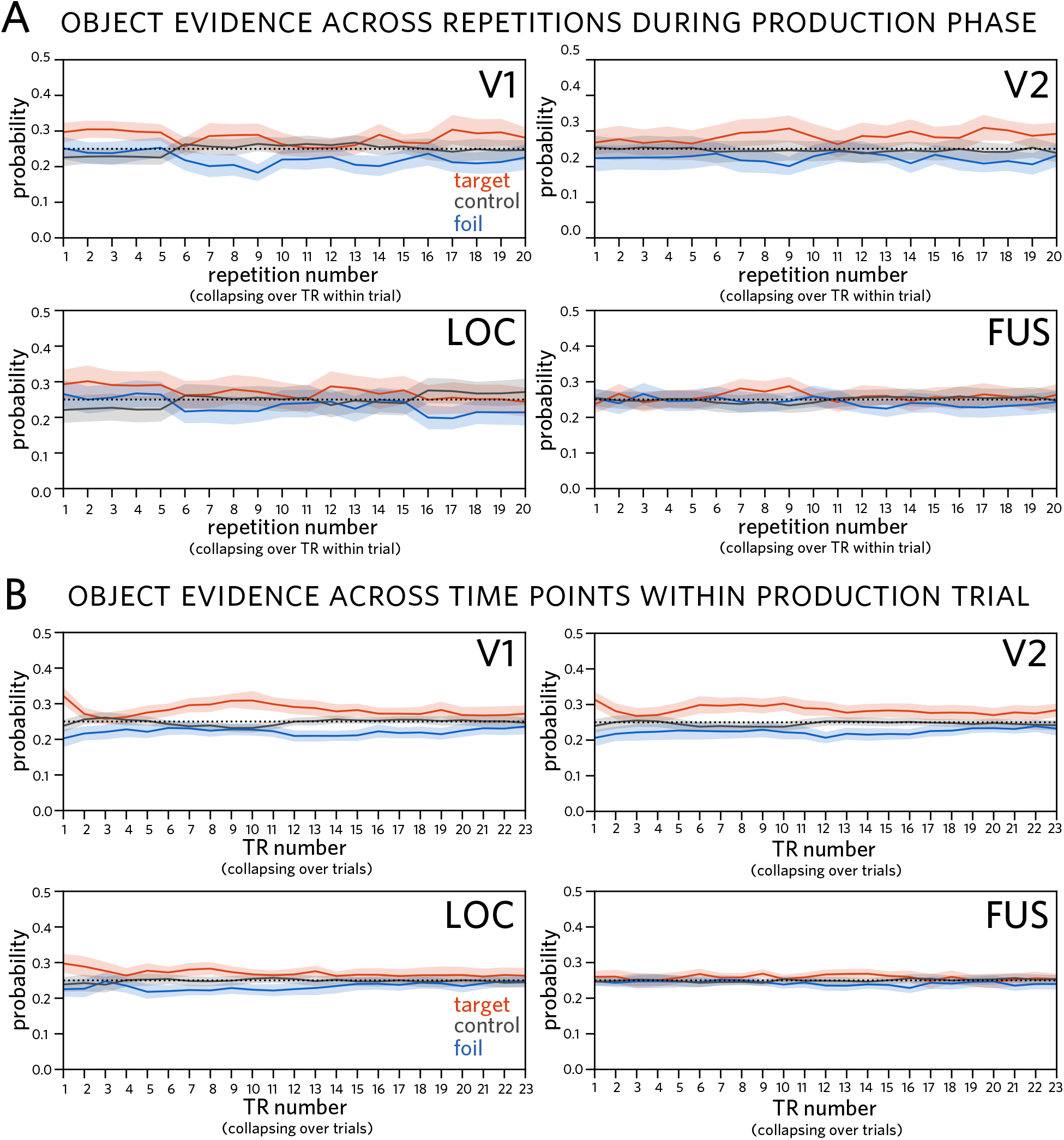
Classifier evidence for each object over time during production, trained on recognition activation patterns. (A) Classifier evidence for target (currently drawn), foil (other trained), and control (never drawn) objects across repetitions during production phase in V1, V2, LOC, and FUS, averaging over TR within trial. (B) Classifier evidence for each object by TR within trial in the same regions, averaging over trials. Probabilities assigned by a 4-way logistic regression classifier trained on patterns of neural responses evoked during initial recognition of these objects. Shaded regions reflect 95% confidence bands.

### Stable object representations in activation patterns in visual cortex

Because we collected neural responses to each object both before and after the production phase, we could also evaluate the consequences of repeatedly drawing an object on the discriminability of neural activation patterns associated with each object in these regions. Insofar as repeatedly drawing the trained objects led to more discriminable representations of those objects *within* each region, we hypothesized that trained object representations would become more differentiated following training, resulting in enhanced object decoding accuracy in the post-production phase relative to the pre-production phase, especially for trained objects. To test this hypothesis, we analyzed changes using a linear mixed-effects model with phase (pre vs. post) and condition (trained vs. control) as predictors of decoding accuracy, with random intercepts for each participant. We did not find evidence that objects differed in discriminability between the pre- and post-production recognition phases in any ROI (i.e., no main effect of phase; *p*s > .225), nor evidence for larger change in discriminability for trained vs. control objects (i.e., no phase by condition interaction *p*s > .135). These results suggest that stimulus-evoked neural activation patterns in occipital cortex were stable under the current manipulation of visual production experience.

### Enhanced object evidence in connectivity patterns among occipitotemporal and parietal regions

Drawing is a complex visuomotor behavior, involving the concurrent recruitment of both occipitotemporal and parietal lobes (Vinci-Booher et al., 2018). In agreement with prior work, we found consistent engagement in voxels within V1, V2, LOC, and parietal cortex during drawing production relative to rest, as measured by a univariate contrast (see Methods). We asked whether this joint engagement may reflect, at least in part, the transmission of object-specific information among these regions. If so, then learning to draw an object across repeated attempts may lead to enhanced transmission of the diagnostic features of the object. To explore whether there was enhanced transmission of object-specific information, we investigated the *connectivity patterns* between voxels in V1, V2, LOC, and parietal cortex engaged during the production task.

Specifically, we computed voxel-wise connectivity matrices between each pair of these ROIs across the duration of each 23-TR trial. We then trained a 2-way logistic regression classifier to predict object identity from these features. For each trial, this classifier yielded two probability values corresponding to the amount of evidence for the *target* and *foil* objects. As in the previous analysis, we computed a target selection log odds ratio, this time for each trial and for each participant. The trials were then divided based on whether they were early (runs 1 and 2) or late (runs 3 and 4) in the production phase. We then analyzed changes in target selection as a function of time using a linear mixed-effects model with random intercepts for each participant.

When analyzing changes in connectivity patterns between V1 and V2, including time as a factor improved model fit, χ^*2*^(1) = 9.078, *p* = 0.0026, β_*time*_ = 0.473, 95% CI = [0.208 0.769]. We found a similar pattern of results for V1/LOC (χ^*2*^(1) = 9.301, *p* = 0.0023, β_*time*_ = 0.456, 95% CI = [0.166 0.720]), for V1/parietal (χ^*2*^(1) = 7.254, *p* = 0.0071, β_*time*_ = 0.409; 95% CI = [0.078 0.723]), for V2/LOC (χ^*2*^(1) = 6.775, *p* = 0.0092, β_*time*_ = 0.388; 95% CI = [0.073 0.701]), and modestly for V2/Parietal (χ^*2*^(1) = 4.293, *p* = 0.038, β_*time*_ = 0.304; 95% CI = [0.024 0.580]) We also analyzed changes in the connectivity pattern for LOC/Parietal, but found no evidence for reliable changes across time (χ^*2*^(1) = 1.01, *p* = 0.3151, β_*time*_ = 0.141; 95% CI = [-0.152 0.407]).

Overall, these results suggest that repeated drawing practice may lead to enhanced transmission of object-specific information among regions in occipital and parietal cortex over time (i.e. from early to late in production phase, Fig. 6).

**Figure 6:**
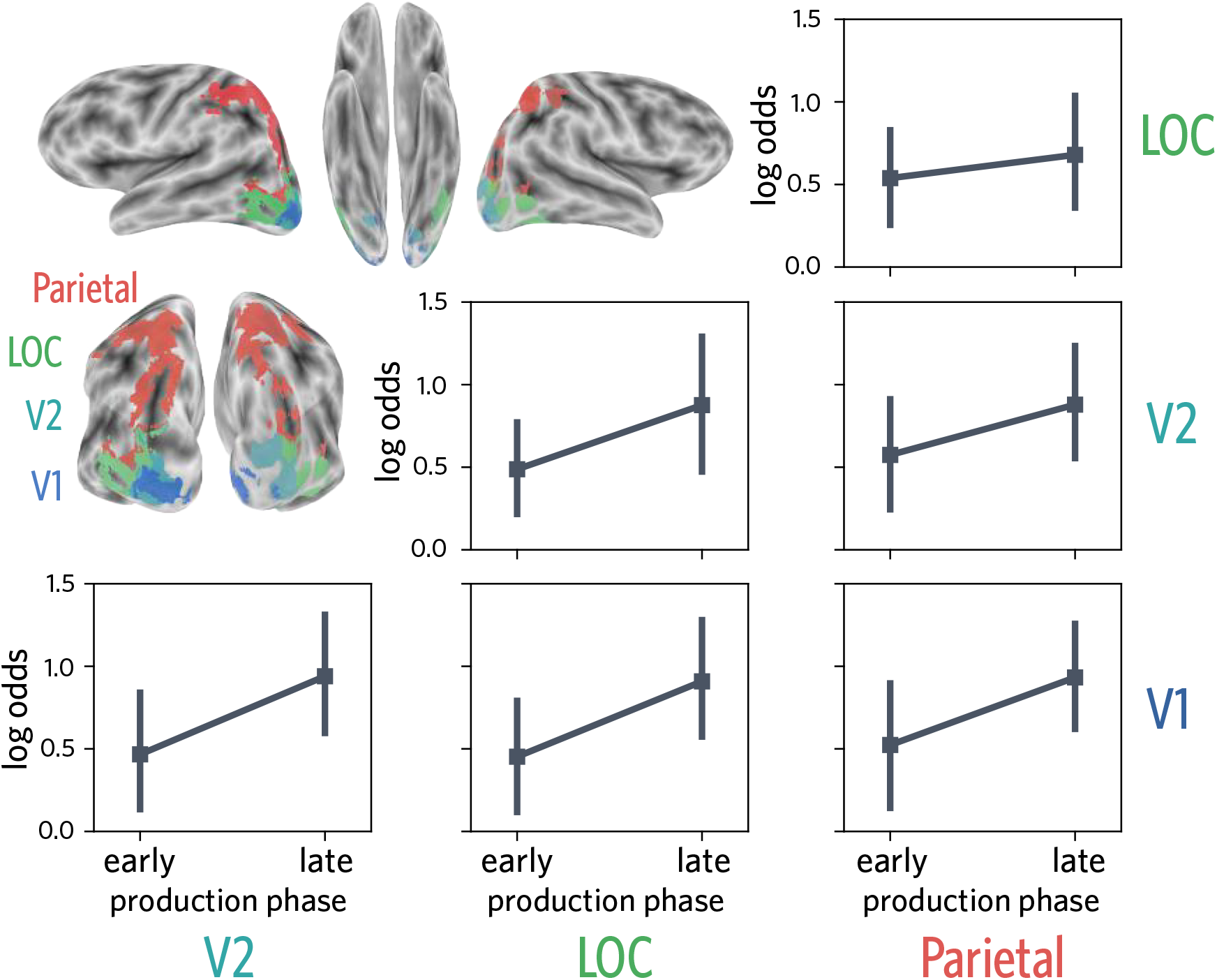
Target selection over time during production, trained on connectivity patterns between pairs of regions. Target selection measured using connectivity patterns in drawing-relevant ROIs taken from V1, V2, LOC and parietal cortex (upper left). Target selection increased from early (runs 1 and 2) to late (runs 3 and 4) phases between V1 and V2, V1 and LOC, and V1 and Parietal.

### Enhanced object evidence not found in activation patterns within regions

To determine whether connectivity patterns between ROIs carried task-related information that was not redundant with information directly accessible (i.e., linearly decodable) from activation patterns within regions, we constructed the same type of classifier on the concatenated voxel activation patterns extracted from each ROI, rather than their connectivity patterns. By contrast with decoding from connectivity patterns, we found that when using concatenated activation patterns from V1 and V2, including time as a factor did not improve the model, χ^*2*^(1) = 0.075, *p* = 0.784, and time did not predict target selection, β_*time*_ = −0.030, 95% CI = [-0.257 0.181]). There were similarly null effects for concatenated V1/LOC (χ^*2*^(1) = 0.690, *p* = 0.406, β_*time*_ = 0.092, 95% CI = −0.126 0.302), V1/Parietal (χ^*2*^(1)=0.203, *p* = 0.652, β_*time*_ = −0.054, 95% CI = [-0.315 0.180]), V2/LOC (χ^*2*^(1) = 0.274, *p* = 0.601, β_*time*_ = 0.059, 95% CI = [-0.171 0.273]), V2/Parietal (χ^*2*^(1) = 0.301, *p* = 0.583, β_*time*_ = −0.066, 95% CI = [-0.315 0.158]), and LOC/Parietal (χ^*2*^(1) = 0.000, *p* = 0.988, β_*time*_ = −0.002, 95% CI = [-0.246 0.251]).

Taken together, these results suggest that connectivity patterns between regions *selectively* carried information about the increasing discriminability of trained object representations over the course of the visual production phase. Such enhanced transmission may reflect participants’ increasing ability to emphasize the diagnostic features of each object across repeated attempts to transform their perceptual representation of the object into an effective motor plan to draw it. More broadly, these findings sugges that the manner in which information is communicated *between* sensory and downstream regions may be a potential neural substrate for learning complex visually guided actions, including visual production (Vinci-Booher et al., 2016).

## Discussion

The current study investigated the functional relationship between recognition and production of objects in human visual cortex. Moreover, we aimed to characterize the consequences of repeated production on the discriminability of object representations. To this end, we scanned participants using fMRI while they performed both recognition and production of the same set of objects. During the production task, they repeatedly produced drawings of two objects. During the recognition task, they repeatedly discriminated the repeatedly drawn objects, as well as a pair of other control objects. We measured spatial patterns of voxel activations in ventral visual stream during drawing production and found that regions in occipital cortex carried diagnostic information about the identity of the currently drawn object that was similar in format to the pattern evoked during visual recognition of a realistic rendering of that object. Moreover, we found that these production-related activation patterns reflected sustained prioritization of the currently drawn object in visual cortex and concurrent suppression of the other repeatedly drawn object, suggesting that visual production recruits an internal representation of the current object to be drawn that emphasizes its diagnostic features. Finally, we found that patterns of functional connectivity between voxels in occipital cortex and parietal cortex supported progressively better decoding of the currently drawn object across the production phase, suggesting a potential neural substrate for production-related learning. Taken together, these findings contribute to our understanding of the neural mechanisms underlying complex behaviors that require the engagement of and interaction between regions supporting perception and action in the brain.

Our findings advance an emerging literature on the neural correlates of visually cued drawing behavior. The studies that comprise this literature have employed widely varying protocols for cueing and collecting drawing data. For example, one early study briefly presented watercolor images of objects as visual cues, and instructed participants to use their right index finger, which lay by their side and out of view, to ‘draw’ the object in the air (Makuuchi, Kaminaga, & Sugishita, 2003). Another study used cartoon images of faces and had participants produce their drawings on a paper-based drawing pad, also hidden from view (Miall, Gowen, & Tchalenko, 2009). More recently however, MR-compatible digital tablets have enabled researchers to automatically capture natural drawing behavior in a digital format while participants are concurrently scanned using fMRI. In one such study, participants copied geometric patterns (i.e., spiral, zigzag, serpentine), which were then projected onto a separate digital display (Yuan & Brown, 2014), while another had participants copy line drawings of basic objects, but participants were unable to view the strokes they had created (Planton, Longcamp, Péran, Demonet, & Jucla, 2017).

Unlike the way people produce drawings in everyday life, participants in these studies generally did not receive visual feedback about the perceptual properties of their drawing while producing it (cf. Yuan & Brown, 2014), and were cued to produce simple abstract shapes rather than real-world objects. By contrast, in the current study we employed photorealistic renderings of 3D objects as visual cues and gave participants continuous visual access to their drawing while producing it. Using photorealistic object stimuli rather than geometric patterns or pre-existing line drawings of objects allowed us to interrogate the functional relationship between the perceptual representations formed during visual recognition of real-world objects and those that are recruited online to facilitate drawing production. Moreover, participants in our study received immediate visual feedback about the perceptual properties of their drawing while producing it, allowing us to investigate distinctive aspects of *drawing* behavior that are not shared with other depictive actions (e.g., gesture) that do not leave persistent visible traces.

Previous studies were also primarily concerned with characterizing overall differences in BOLD signal amplitude between a visually cued drawing and another baseline visual task (i.e., object naming, subtraction of two visually presented numbers). The current study diverges from prior work in its use of machine learning techniques to analyze the expression of object-diagnostic information within visual cortex, as well as in the pattern of connections to downstream parietal regions. As a consequence, our study helps to elucidate the neural content and circuitry that underlie visual production behavior.

The current findings are generally consistent with prior work in observing broad recruitment of a network of regions during visually guided drawing production, including regions in the ventral stream and in parietal regions. Moreover, our findings are congruent with a growing body of evidence showing a large degree of functional overlap in the network of regions during the perception and production of abstract symbols (James, 2017; James & Gauthier, 2006), as well as overlap between regions recruited during production of symbols and object drawings (Planton et al., 2017). This convergence suggests that common functional principles (Lake, Salakhutdinov, & Tenenbaum, 2015), if not identical neural mechanisms, may underlie fluent perception and production of symbols and drawable objects, in particular the recruitment of representations in visual cortex and computations in parietal cortex that are thought to transform perceptual representations into actions (Vinci-Booher et al., 2018).

Interestingly, the most robust information about which object participants were currently drawing was available in occipital cortex. These results are largely consistent with prior work that has found functional overlap between neural representations of perceptual information and information in visual working memory (Sprague, Ester, & Serences, 2014; Harrison & Tong, 2009) and visual imagery (Dijkstra, Bosch, & van Gerven, 2017; Kosslyn, Ganis, & Thompson, 2001). Thus a natural implication for our understanding of the neural mechanisms underlying visual production is that mechanisms supporting visual working memory and visual imagery in sensory cortex are also recruited during production of a drawing of an object held in working memory. And these mechanisms may have provided the basis for our ability to decode the identity of the target object during drawing production.

Although we did not find that trained object representations became more differentiated within our ROIs, we discovered in exploratory analyses that the pattern of connectivity between visual cortex and parietal regions during drawing production carried increasingly diagnostic information about the target object across the production phase. This finding suggests while activation patterns evoked by objects within occipital cortex may not differentiate as a result of repeated production, that the manner in which this information is transmitted to downstream action-oriented regions might. For instance, some visual properties may map selectively onto specific motor plans, such that otherwise similar stimuli may lead to increasingly different actions as participants learn to emphasize the visual properties of an object that distinguish it from other objects in their drawings. Indeed, one possibility raised by this finding is that such a mechanism may underlie the enhanced perceptual discrimination behavior measured in prior work investigating the consequences of repeated visual production (Fan et al., 2018). While the current study was focused on learning-related consequences of visual production practice, other learning studies that have employed different tasks, including categorization training (Jiang et al., 2007), associative memory retrieval (Favila, Chanales, & Kuhl, 2016), spatial route learning (Chanales, Oza, Favila, & Kuhl, 2017), and statistical learning (Schapiro et al., 2012), have found that repeated engagement with similar items can lead to their differentiation in the brain.

Taken together, our findings contribute support for the notion that one route by which learning may occur during visual production is by enhancing the discriminability between the neural representations of repeatedly practiced items, and these representations may be measured as the pattern of activations across voxels within a region, as well as pattern of connectivity between voxels between regions (Wang, Cohen, Li, & Turk-Browne, 2015). In the long run, further application of such multivariate analysis approaches to neural data collected during visual production may shed new light not only on the representation of task-relevant information in sensory cortex, but also how this information is transmitted to downstream parietal and frontal regions that support the planning and execution of complex motor plans (James, 2017; Goodale & Milner, 1992).

## Code accessibility

The code for the analyses presented in this article will be made publicly available in a Github repository upon acceptance of this manuscript.

## Data accessibility

The data presented in this article will be made publicly available upon acceptance of this manuscript.

## Acknowledgments

This work was supported by NSF GRFP DGE-0646086 to J.E.F., NIH R01 EY021755, R01 MH069456, and the David A. Gardner ’69 Magic Project at Princeton University. Thanks to the Computational Memory and Turk-Browne labs for helpful comments.

## Author contributions statement

J.E.F., D.L.K.Y., N.B.T.-B., K.A.N. designed the study. J.E.F. performed the experiments. J.E.F., J.D.W., and J.B.G. conducted analyses. J.E.F., J.D.W., J.B.G., K.A.N., and N.B.T.-B. planned analyses, interpreted results, and wrote the paper.

## References

Bainbridge, W. A., Hall, E. H., & Baker, C. I. (2019). Drawings of real-world scenes during free recall reveal detailed object and spatial information in memory. Nature communications, 10(1), 5.

Bates, D., Maechler, M., Bolker, B., Walker, S., Christensen, R. H. B., Singmann, H., … Rcpp, L. (2015). Package lme4. Convergence, 12(1).

Chanales, A. J., Oza, A., Favila, S. E., & Kuhl, B. A. (2017). Overlap among spatial memories triggers repulsion of hippocampal representations. Current Biology, 27(15), 2307–2317.

Chun, M. M., Golomb, J. D., & Turk-Browne, N. B. (2011). A taxonomy of external and internal attention. Annual review of psychology, 62, 73–101.

Cichy, R. M., Chen, Y., & Haynes, J.-D. (2011). Encoding the identity and location of objects in human loc. Neuroimage, 54(3), 2297–2307.

Cohen, J. D., Daw, N., Engelhardt, B., Hasson, U., Li, K., Niv, Y., … others (2017). Computational approaches to fmri analysis. Nature Neuroscience, 20(3), 304.

Davachi, L. (2006). Item, context and relational episodic encoding in humans. Current opinion in neurobiology, 16(6), 693–700.

Dijkstra, N., Bosch, S. E., & van Gerven, M. A. (2017). Vividness of visual imagery depends on the neural overlap with perception in visual areas. Journal of Neuroscience, 37(5), 1367–1373.

Draschkow, D., Wolfe, J. M., & Vo, M. L.-H. (2014). Seek and you shall remember: Scene semantics interact with visual search to build better memories. Journal of Vision, 14(8), 10–10.

Epstein, R., Graham, K. S., & Downing, P. E. (2003). specific scene representations in human parahippocampal cortex. Neuron, 37(5), 865–876.

Fan, J. E., & Turk-Browne, N. B. (2013). Internal attention to features in visual short-term memory guides object learning. Cognition, 129, 2.

Fan, J. E., Yamins, D. L. K., & Turk-Browne, N. B. (2018). Common object representations for visual production and recognition. Cognitive Science, 0(0). doi: 10.1111/cogs.12676

Favila, S. E., Chanales, A. J., & Kuhl, B. A. (2016). Experience-dependent hippocampal pattern differentiation prevents interference during subsequent learning. Nature communications, 7, 11066.

Garvert, M. M., Dolan, R. J., & Behrens, T. E. (2017). A map of abstract relational knowledge in the human hippocampal–entorhinal cortex. Elife, 6, e17086.

Gazzaley, A., & Nobre, A. C. (2012). Top-down modulation: bridging selective attention and working memory. Trends in cognitive sciences, 16(2), 129–135.

Gegenfurtner, K. R., Kiper, D. C., & Fenstemaker, S. B. (1996). Processing of color, form, and motion in macaque area v2. Visual neuroscience, 13(1), 161–172.

Goodale, M., & Milner, A. (1992). Separate visual pathways for perception and action. Trends in Neuro-sciences, 15(1), 20–25.

Grill-Spector, K., Kourtzi, Z., & Kanwisher, N. (2001). The lateral occipital complex and its role in object recognition. Vision research, 41(10-11), 1409–1422.

Gross, C. G. (1992). Representation of visual stimuli in inferior temporal cortex. Philosophical Transactions of the Royal Society of London. Series B: Biological Sciences, 335(1273), 3–10.

Güçlü, U., & van Gerven, M. A. (2015). Deep neural networks reveal a gradient in the complexity of neural representations across the ventral stream. Journal of Neuroscience, 35(27), 10005–10014.

Harrison, S. A., & Tong, F. (2009). Decoding reveals the contents of visual working memory in early visual areas. Nature, 458(7238), 632.

Haxby, J. V., Gobbini, M. I., Furey, M. L., Ishai, A., Schouten, J. L., & Pietrini, P. (2001). Distributed and overlapping representations of faces and objects in ventral temporal cortex. Science, 293(5539), 2425–2430.

Hubel, D. H., & Wiesel, T. N. (1968). Receptive fields and functional architecture of monkey striate cortex. The Journal of physiology, 195(1), 215–243.

Hung, C. P., Kreiman, G., Poggio, T., & DiCarlo, J. J. (2005). Fast readout of object identity from macaque inferior temporal cortex. Science, 310(5749), 863–866.

James, K. H. (2017). The importance of handwriting experience on the development of the literate brain. Current Directions in Psychological Science, 26(6), 502–508.

James, K. H., & Atwood, T. P. (2009). The role of sensorimotor learning in the perception of letter-like forms: Tracking the causes of neural specialization for letters. Cognitive Neuropsychology, 26(1), 91–110.

James, K. H., & Gauthier, I. (2006). Letter processing automatically recruits a sensory–motor brain network. Neuropsychologia, 44(14), 2937–2949.

Jiang, X., Bradley, E., Rini, R. A., Zeffiro, T., VanMeter, J., & Riesenhuber, M. (2007). Categorization training results in shape- and category-selective human neural plasticity. Neuron, 53(6), 891–903.

Kamitani, Y., & Tong, F. (2005). Decoding the visual and subjective contents of the human brain. Nature neuroscience, 8(5), 679.

Kosslyn, S. M., Ganis, G., & Thompson, W. L. (2001). Neural foundations of imagery. Nature reviews neuroscience, 2(9), 635.

Kourtzi, Z., & Kanwisher, N. (2001). Representation of perceived object shape by the human lateral occipital complex. Science, 293(5534), 1506–1509.

Lake, B. M., Salakhutdinov, R., & Tenenbaum, J. B. (2015). Human-level concept learning through probabilistic program induction. Science, 350(6266), 1332–1338.

Lewis-Peacock, J. A., & Postle, B. R. (2012). Decoding the internal focus of attention. Neuropsychologia, 50(4), 470–478.

Li, J. X., & James, K. H. (2016). Handwriting generates variable visual output to facilitate symbol learning. Journal of Experimental Psychology: General, 145(3), 298.

Longcamp, M., Boucard, C., Gilhodes, J.-C., Anton, J.-L., Roth, M., Nazarian, B., & Velay, J.-L. (2008). Learning through hand- or typewriting influences visual recognition of new graphic shapes: Behavioral and functional imaging evidence. Journal of cognitive neuroscience, 20(5), 802–815.

Makuuchi, M., Kaminaga, T., & Sugishita, M. (2003). Both parietal lobes are involved in drawing: a functional mri study and implications for constructional apraxia. Cognitive brain research, 16(3), 338–347.

Miall, R. C., Gowen, E., & Tchalenko, J. (2009). Drawing cartoon faces–a functional imaging study of the cognitive neuroscience of drawing. Cortex, 45(3), 394–406.

Murray, E. A., & Bussey, T. J. (1999). Perceptual–mnemonic functions of the perirhinal cortex. Trends in cognitive sciences, 3(4), 142–151.

Norman, K. A., Polyn, S. M., Detre, G. J., & Haxby, J. V. (2006). Beyond mind-reading: multi-voxel pattern analysis of fmri data. Trends in cognitive sciences, 10(9), 424–430.

Peynircioğlu, Z. F. (1989). The generation effect with pictures and nonsense figures. Acta Psychologica, 70(2), 153–160.

Planton, S., Longcamp, M., Péran, P., Demonet, J.-F., & Jucla, M. (2017). How specialized are writing-specific brain regions? an fmri study of writing, drawing and oral spelling. Cortex, 88, 66–80.

Rust, N. C., & DiCarlo, J. J. (2010). Selectivity and tolerance (invariance) both increase as visual information propagates from cortical area v4 to it. The Journal of Neuroscience, 30(39), 12978–12995.

Sayim, B., & Cavanagh, P. (2011). What line drawings reveal about the visual brain. Frontiers in human neuroscience, 5.

Schapiro, A. C., Kustner, L. V., & Turk-Browne, N. B. (2012). Shaping of object representations in the human medial temporal lobe based on temporal regularities. Current Biology, 22(17), 1622–1627.

Serences, J. T., Ester, E. F., Vogel, E. K., & Awh, E. (2009). Stimulus-specific delay activity in human primary visual cortex. Psychological science, 20(2), 207–214.

Sprague, T. C., Ester, E. F., & Serences, J. T. (2014). Reconstructions of information in visual spatial working memory degrade with memory load. Current Biology, 24(18), 2174–2180.

Tipper, S. P., Weaver, B., & Houghton, G. (1994). Behavioural goals determine inhibitory mechanisms of selective attention. The Quarterly Journal of Experimental Psychology Section A, 47(4), 809–840.

Vinci-Booher, S., Cheng, H., & James, K. H. (2018). An analysis of the brain systems involved with producing letters by hand. Journal of cognitive neuroscience, 1–17.

Vinci-Booher, S., James, T. W., & James, K. H. (2016). Visual-motor functional connectivity in preschool children emerges after handwriting experience. Trends in Neuroscience and Education, 5(3), 107–120.

Walther, D. B., Chai, B., Caddigan, E., Beck, D. M., & Li, F. F. (2011). Simple line drawings suffice for functional mri decoding of natural scene categories. Proceedings of the National Academy of Sciences, 108(23), 9661–9666.

Wammes, J. D., Meade, M. E., & Fernandes, M. A. (2016). The drawing effect: Evidence for reliable and robust memory benefits in free recall. The Quarterly Journal of Experimental Psychology, 69(9), 1752–1776.

Wang, Y., Cohen, J. D., Li, K., & Turk-Browne, N. B. (2015). Full correlation matrix analysis (fcma): An unbiased method for task-related functional connectivity. Journal of neuroscience methods, 251, 108–119.

Yamins, D. L., Hong, H., Cadieu, C. F., Solomon, E. A., Seibert, D., & DiCarlo, J. J. (2014). Performance-optimized hierarchical models predict neural responses in higher visual cortex. Proceedings of the National Academy of Sciences, 201403112.

Yuan, Y., & Brown, S. (2014). The neural basis of mark making: A functional mri study of drawing. PloS one, 9(10), e108628.

